# SmMIP-tools: a computational toolset for processing and analysis of single-molecule molecular inversion probes derived data

**DOI:** 10.1101/2021.06.03.446993

**Authors:** Jessie J. F. Medeiros, Jose-Mario Capo-Chichi, Liran I. Shlush, John E. Dick, Andrea Arruda, Mark D. Minden, Sagi Abelson

## Abstract

Single-molecule molecular inversion probes (smMIPs) provides a modular and cost-effective platform for high-multiplex targeted next-generation sequencing (NGS). Nevertheless, translating the raw smMIP-derived sequencing data into accurate and meaningful information currently requires proficient computational skills and a large amount of computational work, prohibiting wide-scale adoption of smMIP-based technologies. To enable easy, efficient, and accurate interrogation of smMIP-derived data, we developed SmMIP-tools, a computational toolset that combines the critical analytic steps for smMIP data interpretation into a single computational pipeline. Here, we describe in detail two of the software’s major components. The first is a read processing tool that performs quality control steps, generates read-smMIP linkages and retrieves molecular tags. The second is an error-aware variant caller capable of detecting single nucleotide variants (SNVs) and short insertions and deletions (indels). Using a cell-line DNA dilution series and a cohort of blood cancer patients, we benchmarked SmMIP-tools and evaluated its performance against clinical sequencing reports. We anticipate that SmMIP-tools will increase accessibility to smMIP-technology, enabling cost-effective genetic research to push personalized medicine forward.

## Introduction

Padlock probes, initially described in 1994 (Nilsson et al. 1994), are linear DNA oligos that contain two target arm segments, physically connected by a linker sequence, designed to complement immediately adjacent target DNA. Upon hybridization of both complementary arms to their target, padlock probes can circularize via ligation, capturing the DNA molecule in a “padlock” conformation to enable localized detection of DNA. The concept of padlock probes was later extended via the introduction of molecular inversion probe (MIP) technology for genotyping of single nucleotide polymorphisms (SNP) (Hardenbol et al. 2003, 2005). Here, probes were designed such that target arms contained a single nucleotide gap flanking a SNP of interest. Upon MIP hybridization, gap-filling by DNA polymerase with the correct deoxynucleoside triphosphate allowed the capture of the SNP locus, later amplified by PCR with standard primers and read-out using DNA microarray. Critically, these studies exhibit unprecedented levels of MIP multiplex capabilities by combining thousands of MIPs in a single reaction while maintaining high levels of specificity. This is in contrast to the notoriously poor multiplex capabilities of standard PCR (Mamanova et al. 2010). Thus, specificity and multiplex capability have positioned MIPs as a vital tool for target enrichment in DNA sequencing. Indeed, one study utilizing MIPs designed with larger gaps between target arms for the multiplex capture and amplification of intervening sequence showed that thousands of exons could be interrogated in a single reaction with subsequent shotgun sequencing (Porreca et al. 2007). However, while individual MIPs were highly specific, uniformity in coverage across MIPs was extremely poor, presenting a major technical challenge. High coverage variability has been addressed with optimized protocols, empirical probe rebalancing, and reduced panel size to enhance uniformity across MIPs (Turner et al. 2009). Critically, these subsequent studies enhanced MIP design elements to permit direct resequencing using NGS platforms such that libraries could be multiplexed to an unprecedented level using barcoded PCR primers for library amplification (O’Roak et al. 2012). High customizability of MIP design has enabled initial groups to generate custom target panels for mutation analysis on large human cohorts, exhibiting the usefulness of MIPs to provide significant biological findings in a high-throughput, cost-effective manner (O’Roak et al. 2012). MIPs continued to evolve and were further enhanced to include unique molecular identifiers (UMIs) into the oligo structure (Hiatt et al. 2013), transforming MIPs into smMIPs and allowing the precise identification of single molecules. UMIs permit the collapse of PCR amplified capture products into single-strand consensus sequences (SSCS), enabling the correction of PCR amplification-induced errors. A second UMI sequence adjacent to a target arm further enhanced smMIP SSCS capabilities (Acuna-Hidalgo et al. 2017).

Though these advancements in smMIP technology have largely addressed the technical challenges of smMIP-based sequencing, there is much room for informatic approaches to enhance capabilities further. This is exemplified by MIPgen (Boyle et al. 2014), a machine learning algorithm that has dramatically improved smMIP panel design, increasing the chances of successful target capture and coverage uniformity across smMIPs. The simplification of panel design has driven the recent uptake of smMIP-based technology for customized, targeted sequencing and subsequent SNV, indel, and in some cases, copy number variant mutational analysis (Neveling et al. 2017; Pippucci et al. 2019; Khan et al. 2020). However, while smMIP-sequencing has been used to generate massive amounts of data, most groups have relied on in-house custom pipelines and there remains a paucity of publicly available informatic tools specifically designed for the effective and efficient handling of smMIP-derived sequencing data. For this reason, we developed SmMIP-tools. SmMIP-tools functionalities include quality control metrics of aligned reads, filtering reads which are inadequate for subsequent analysis, generating base call summaries for both raw reads and SSCS, and the calling of both SNV and short indels with high accuracy. To mitigate false-positive calls arising due to the high level of NGS associated errors (Ma et al. 2019), SmMIP-tools generates probabilistic models of allele-specific error rates. SmMIP-tools dramatically enhances the fidelity of error-corrected mutation calling by analyzing data from each read derived from sequencing read pairs, overlapping smMIPs and smMIP library technical replicates when available. We benchmarked the use of SmMIP-tools via a dilution assay of established cell lines with known mutations. Critically, we validated its utility in side-by-side comparative analysis to detect SNVs and indels previously identified by clinical reports in a large clinical cohort. SmMIP-tools represents the next step in the evolution of smMIP-based technologies as a crucial informatic advance to promote widespread, efficient use of smMIPs to reveal significant biologic findings in a high-throughput and cost-effective manner.

## Results

### Overview

Proper smMIP design (Fig. 1A-C, Methods) is fundamental for the overall success of the sequencing experiment. It is critical to maintain high capture uniformity and specificity and to improve accuracy in detecting rare, subclonal mutations by reducing NGS-associated artefacts. For example, one can improve smMIP panel design by accounting for the sequencing platform intended for use with respect to its output read length. Assuring that each of the paired reads thoroughly interrogates its corresponding target locus will maximize the coverage at each base and support the suppression of errors that preferentially occur on one of the two amplified cDNA strands.

**Figure. 1.**
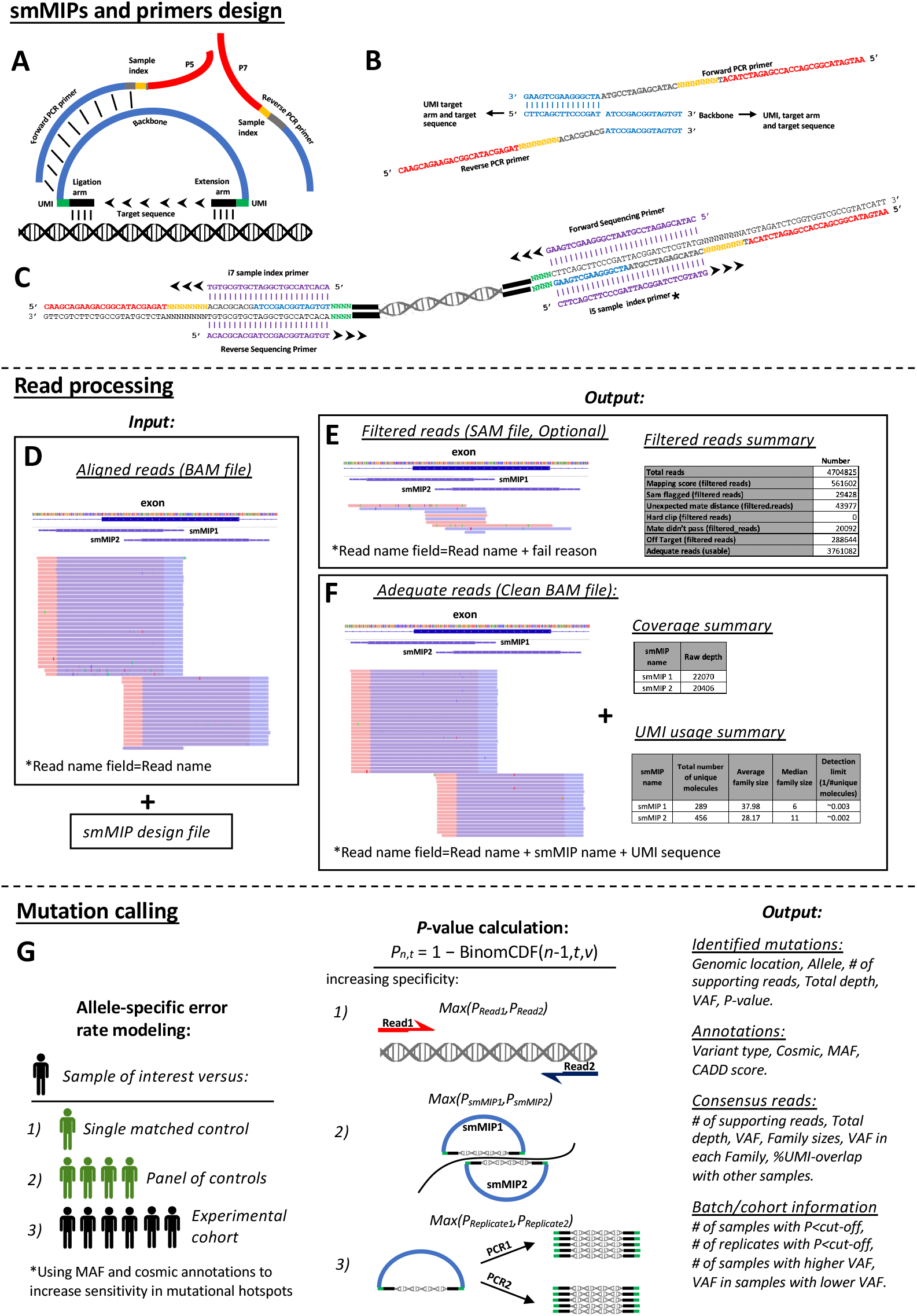
Overview of smMIP assay design and key steps in data analysis using SmMIP-tools. (A) General layout of a smMIP. (B) Backbone and PCR primers’ sequences and primers annealing sites. (C) Library composition and sequencing primers’ sequences. *the i5 sample index primer is only required for select sequencing platforms. >>> indicates the direction of extension. (D) SmMIP-tools BAM processing inputs. (E,F) SmMIP-tools BAM processing outputs. (G) Probabilistic models of error rates and SmMIP-tools mutation calling output. Let ‘v’ be the investigated allele’s fraction in the cohort (option 1,2 or 3 in the left panel). Then, the P-value of seeing at least ‘n’ reads supporting the non-reference allele out of ‘t’ total coverage in the investigated genomic position in the sample of interest is Pn,t = 1 - BinomCDF(n-1,t,v) where BinomCDF denotes the binomial cumulative distribution function.

Following sequencing, quality control and read processing steps are critical to achieving reliable variant discovery (Koboldt 2020). SmMIP-tools takes as an input a read-alignment BAM file and a smMIP design file generated by MIPgen (Boyle et al. 2014) (Fig. 1D). It uses information concerning each probe and its target sequence to apply a set of filters to discard hard-clipped reads, reads with low mapping quality, paired reads with an unexpected insert size or improper alignment orientations (Fig. 1E). To confirm the proper structure of the remaining reads and to identify corrupted UMI sequences, associations between reads and their precise probe-of-origin (termed here, read-smMIP linkages) are generated using two authentication steps. These include the identification of smMIP targets that overlap with the paired reads’ targeted genomic region and validation of a single smMIP among those candidates through local alignment of their extension and ligation arm sequences to the paired reads sequences. The final output contains quality control summary files concerning raw and consensus reads (Fig. 1F) and a BAM file with the remaining high-quality reads. At that point, UMI sequences and smMIP-of-origin are included in each read’s header.

In the following steps, SmMIP-tools uses the processed BAM file to generate base call summaries of raw reads and SSCS which are subsequently those are refined by the software’s error-aware variant detection algorithm (Fig. 1G). To call mutations, binomial models are used to conduct allele-specific frequency comparisons between each sample of interest and either a single control or a cohort of control samples. SmMIP-tools is also capable of precise allele-specific error rates estimation without using dedicated controls by comparing a sample of interest to the remainder of the experimental cohort. Prior knowledge concerning the location of common cancer mutations and germline polymorphisms is used to increase the sensitivity of detecting recurrently mutated alleles (Methods). To improve specificity, non-reference alleles are evaluated separately in each of the paired sequencing reads, in reads derived from different but overlapping smMIPs, and in technical replicates when available. The final output is a comprehensive report that includes the detected mutations, key variant annotations, information concerning consensus reads’ support, sequencing batch summaries and mutations flags; all of which are valuable for the ranking and prioritization of variants.

A comparison matrix emphasizing differences in strengths and limitations between SmMIP-tools and other popular software is included in Supplemental Table 1.

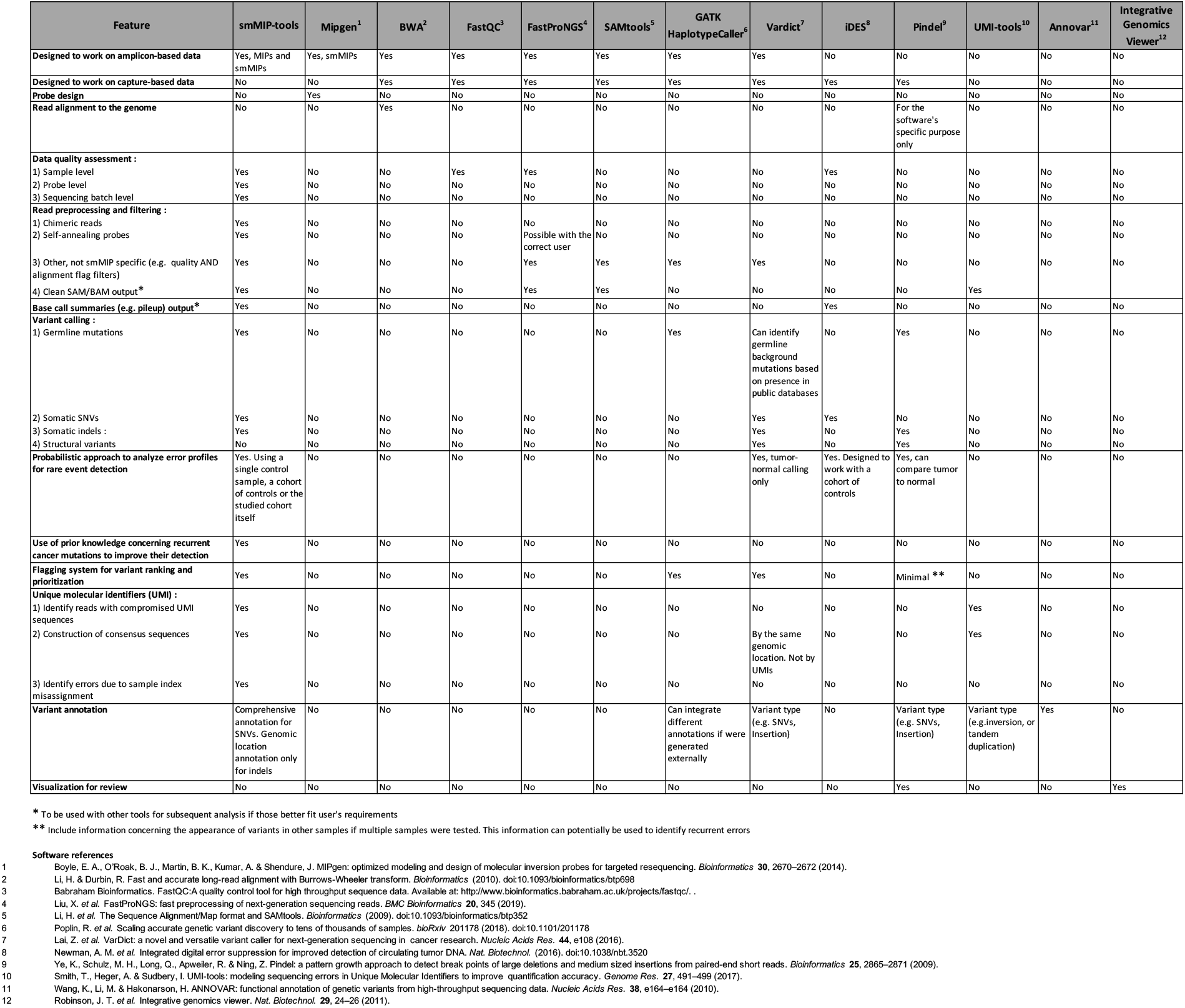

### Evaluating SmMIP-tools ability to generate authentic read-smMIP linkages

Identifying the correct smMIP arm sequences and validating their expected position in the sequenced read pairs is essential to pinpoint self-annealed probes that lack target sequences and to eliminate chimeric inserts generated by partially overlapping probes (Supplemental Fig. 1A). Furthermore, read-smMIP linkages can help to remove mutations that are called outside the target region of individual smMIPs (Supplemental Fig. 1B), to eliminate errors based on ambiguous calls in regions with overlapping smMIPs and to validate the UMIs’ sequence integrity in the expected insert layout (Supplemental Fig. S1C).

**Supplemental Figure 1.**
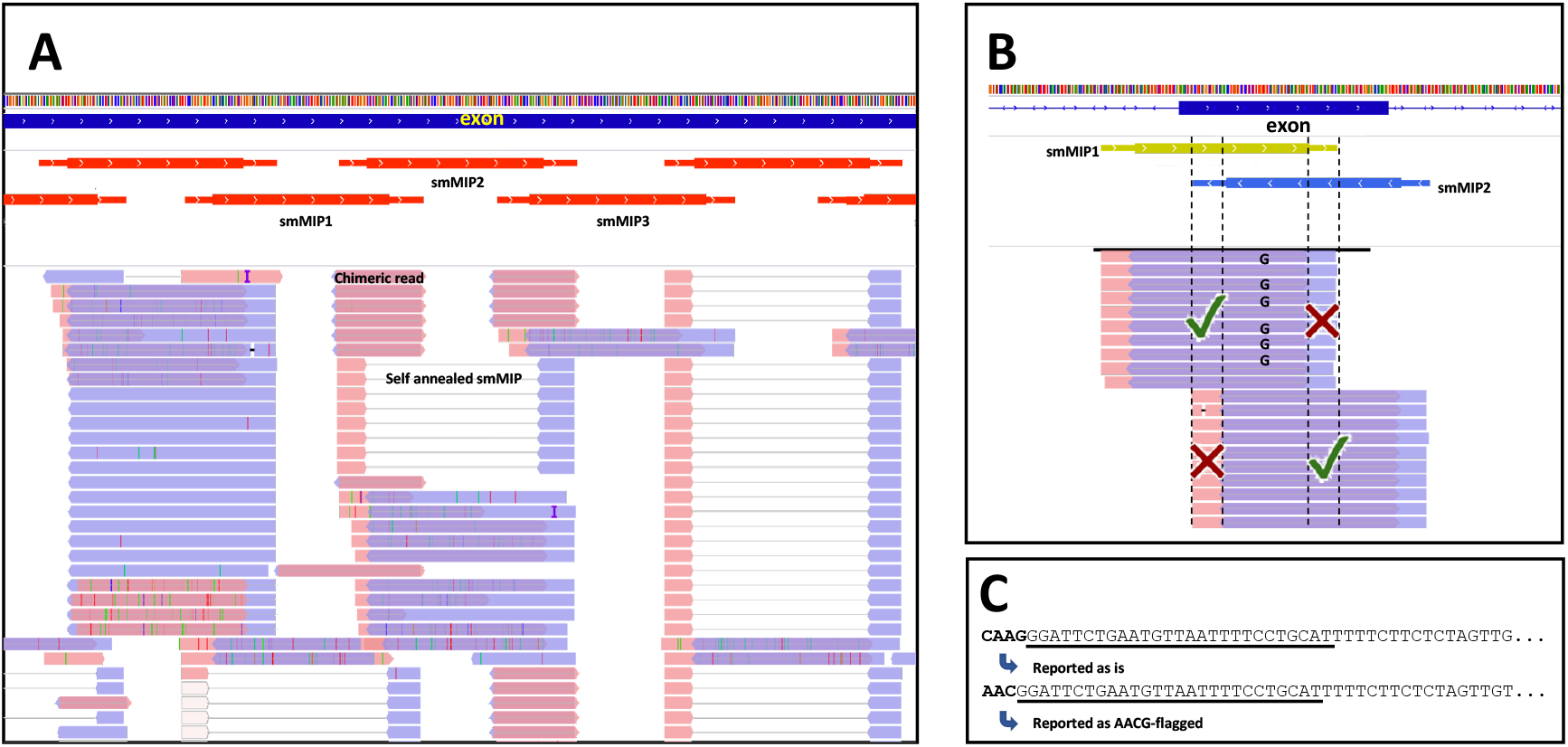
Addressing issues related to the smMIP technology by linking reads to their probe-of-origin with read-smMIP linkages. (A) Read-smMIP linkages can support the identification and removal of typical undesired sequencing library products. The example shows a representative chimeric read generated by one arm that belongs to smMIP1 and another to smMIP2, as well as a read without a target sequence generated from smMIP2 both of which are are emphasised by the added text. (B) Read-smMIP linkages support the identification of the regions targeted by a given smMIP, thus preventing mutation calling from their arm sequences. (C) Validating the correct position of the smMIP arm’s sequences in the expected reads layout helps to identify UMI errors and preserve high quality data for subsequent analysis.

We evaluated the ability of SmMIP-tools to link the correct probe-of-origin to each sequencing read by designing an experiment consisting of real smMIPs that were used for sequencing and simulated smMIPs that were not.

Overall, 22,299 smMIPs were simulated by Mipgen (Boyle et al. 2014), that is, 56-80 simulated smMIPs that overlaps each real smMIP. The simulated probes were designed to include smMIPs with arms and target loci of variable length that can either partially overlap with those of the real smMIPs, are fully included within a real smMIP’s genomic insert, or extend beyond the real smMIPs’ 3’ and/or 5’ direction (Supplemental Fig. 2).

**Supplemental Figure 2.**
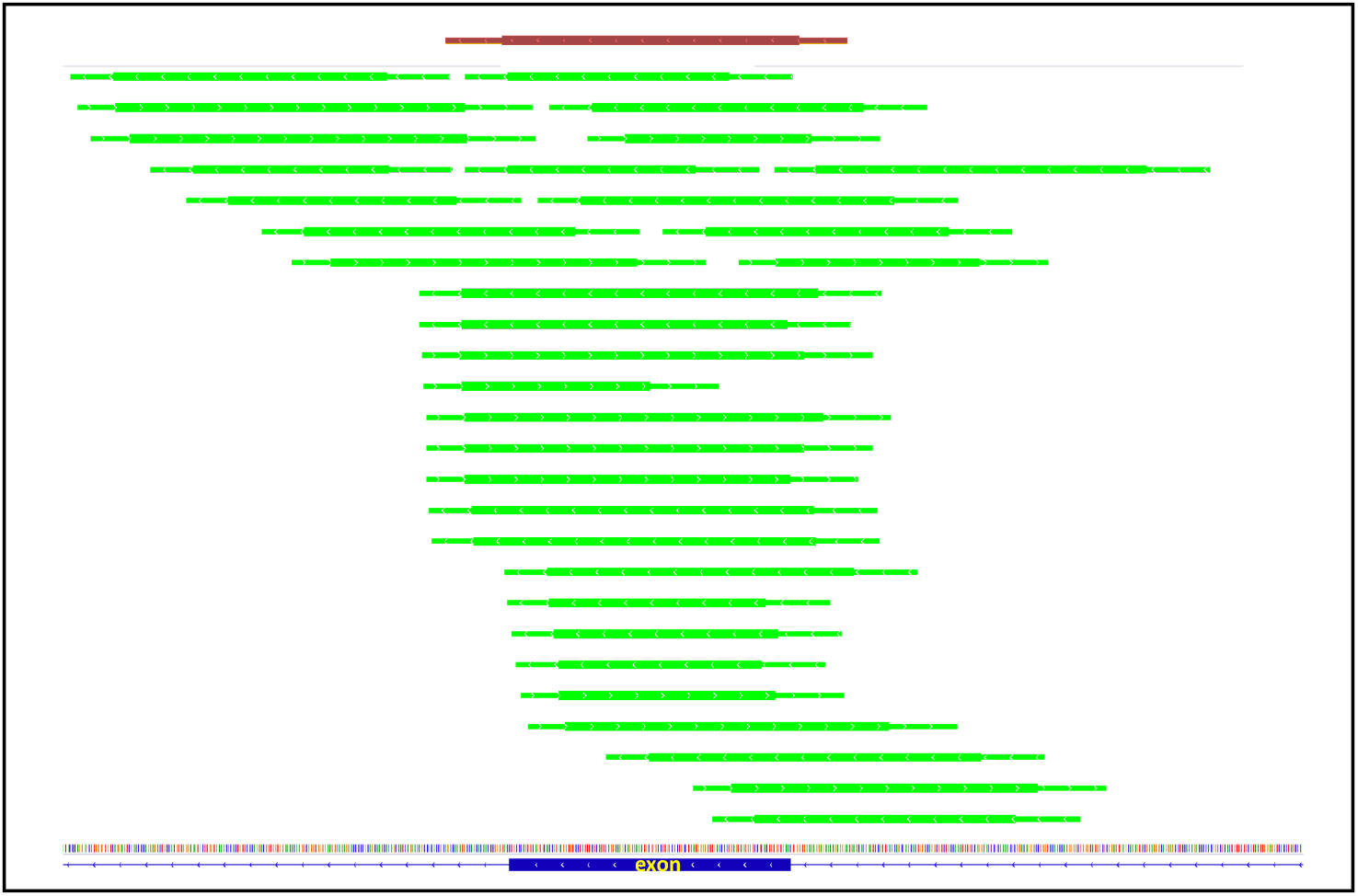
Simulating smMIPs to assess the performance of read-smMIP linkages generation. Simulated smMIPs (green, n=22,299) were designed in the proximity of the real smMIPs that were used for sequencing (brown, n=284). A simulated smMIP can either overlap with a real smMIP, be smaller and included within the genomic loci belonging to a real smMIP or be larger and flank the entire genomic loci belong to a real smMIP. The closer the start and end genomic positions of simulated smMIPs are to those of real smMIPs the harder it is to accurately determine the reads’ origin. Broad and narrow lines represent smMIP’s targeted region and arms respectively.

We further controlled for the distance between real and simulated smMIPs’ ends to separate the simulated smMIPs into six sets (R0-R5). Those sets ranged from R0, which include simulated smMIPs with an identical start or end genomic position as those of real smMIPs (i.e., all the probes are included). Therefore, R0 represents the set of which the task of separating between real and simulated smMIPs is most challenging. On the other extreme is set R5, which in addition to real smMIPs, only includes simulated smMIPs with ends that are more than five genomic bases away from those of the real smMIP. Therefore, with R5 the tasks of accurately assigning the correct probe-of-origin to each read and differentiate between real and simulating smMIPs is expected to be the least challenging.

A total of 25,353,671 read pairs from 16 sequenced cord blood samples were subjected to read-smMIP linkage performance analysis. On average, 6.1% of the reads could not be linked to any smMIP (real or simulated) and were considered “off-target” due to insufficient overlap with the targeted loci (Fig. 2A). From the other 93.9% “on-target” reads, SmMIPs-tools correctly assigned the authentic probe-of-origin to 89.72% of the reads in R0, the most restrictive simulated smMIP set (Fig. 2B). In the most restrictive set, 8.79% of the reads could equally be associated with more than one smMIP, and 0.04% were falsely assigned to a simulated smMIP (Fig. 2C). Performance was significantly improved when simulated smMIPs that start or end at an identical genomic base as real smMIPs were omitted (R1).

**Figure. 2.**
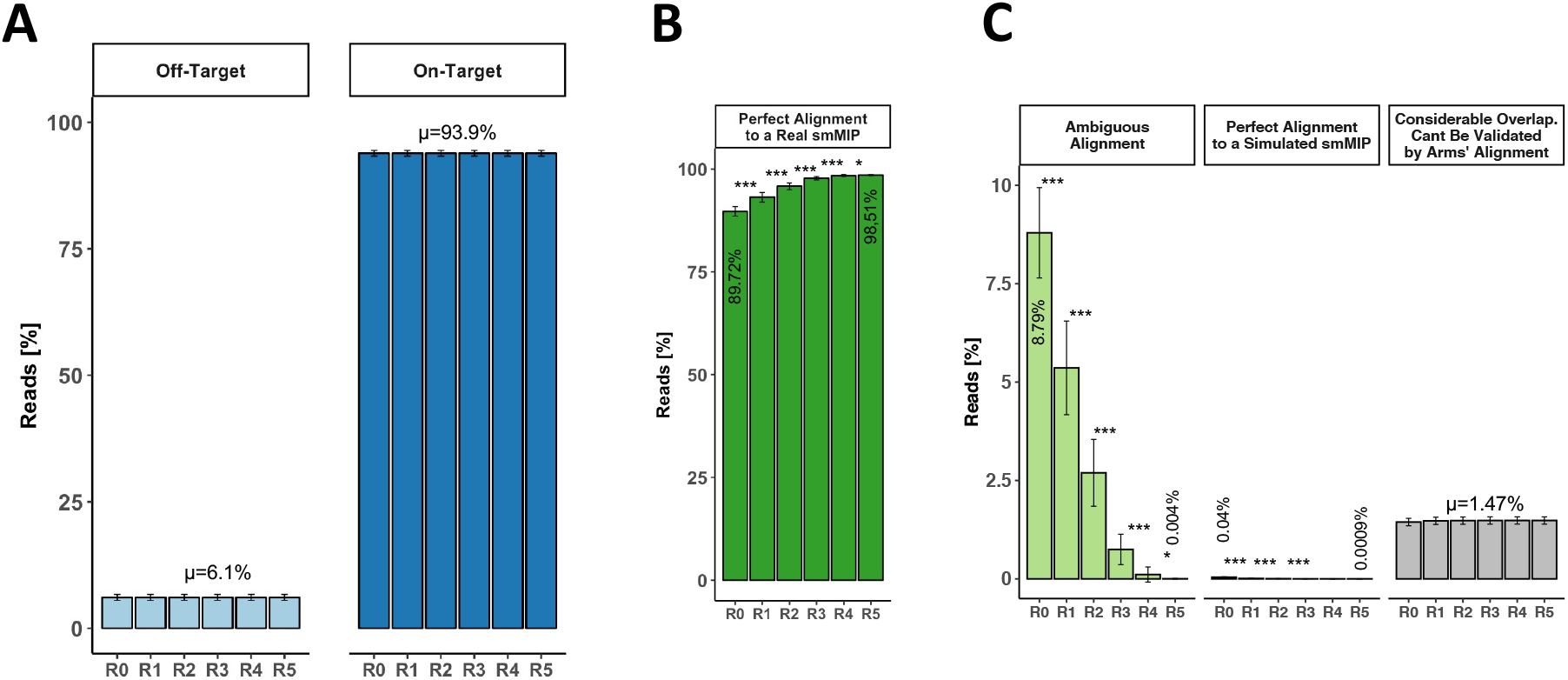
SmMIP-tools delivers accurate read-smMIP linkages allowing proper downstream analysis of data from complex smMIP panels. **(A)** Average percentage of off-target and on-target reads. **(B)** Average percentage of perfect alignment to a real smMIP. **(C)** Average percentage of imperfect alignments including ambiguous alignment, perfect alignment to a simulated smMIP and alignments that could not be validated by target arms. Error bars representing standard deviations for reads derived from 16 sequenced samples. Paired Sample T-Test: * P<0.05, *** P<0.001. Perfect alignment is defined by first searching for smMIPs that their target overlaps with the read’s genome alignment followed by validation of a single smMIP by local alignment of the candidates smMIPs’ arms to the paired reads. R0 includes real smMIPs and simulated smMIPs that start or end at the same genomic location as real smMIPs. R3 for example, includes real smMIPs and simulated smMIPs that start or end more than 3 bases apart from real smMIPs start and end sites. R5 for example corresponds to >5 base difference.

In the most permissive simulated smMIP subset (R5), 98.51% of the on-target reads were correctly assigned to a single real smMIP, and a negligible percentage of the reads were falsely assigned to a simulated smMIP (0.0009%). We noticed that sequencing and library amplification errors confound ambiguous or inaccurate assignment of reads to their probe-of-origin. Specifically, due to non-reference bases, on average, 1.47% of the on-target reads showed considerable overlap with the target locus yet failed validation by local alignment of the probes’ arms to the reads sequence. SmMIP-tools can still salvage such reads. Nevertheless, since their UMI sequence integrity cannot be validated, such reads are flagged and not included in downstream analyses that consider SSCS (Supplemental Fig. 1C). Our results indicate that SmMIP-tools can generate read-smMIP linkages to accurately filter problematic reads and prepare smMIP-derived data from highly complex target panels, including those containing highly overlapping smMIPs for more efficient downstream analyses.

### Benchmarking SmMIP-tools mutations calling algorithm using cell line DNA mixtures

SmMIP-tools incorporates multiple techniques to suppress false positive calls (Fig. 1G). To benchmark their use, we first constructed high confident lists of true and false positive mutations by sequencing 8 leukemia cell lines (Supplemental Table 2, Methods). DNA from each cell line was then mixed to generate 6 separate pools containing varying concentrations of each cell line’s genomic material. Each mix was sequenced twice to enable the use of information derived from technical replicates to enhance error suppression using SmMIP-tools. In each of the 12 sequenced libraries, we counted the number of error-free positions, defined as positions in the interrogated genomic space, represented exclusively with reference alleles before and after applying error suppression techniques. Both the consensus reads assembly and error modelling techniques (Methods) delivered significant levels of error suppression as indicated by the sole presence of reference alleles in 72.83% and 98.47% of the investigated genomic positions respectively; as compared with an average of 1.22% before error correction (Fig. 3A). Error suppression using the binomial error rate modelling approach was further augmented when information derived from separate strands or from separate technical replicates was incorporated (Fig. 3B). Improved error suppression was also observed when alleles that are covered by different overlapping smMIPs were evaluated against error models derived from each smMIP independently (Fig. 3C).

**Figure. 3.**
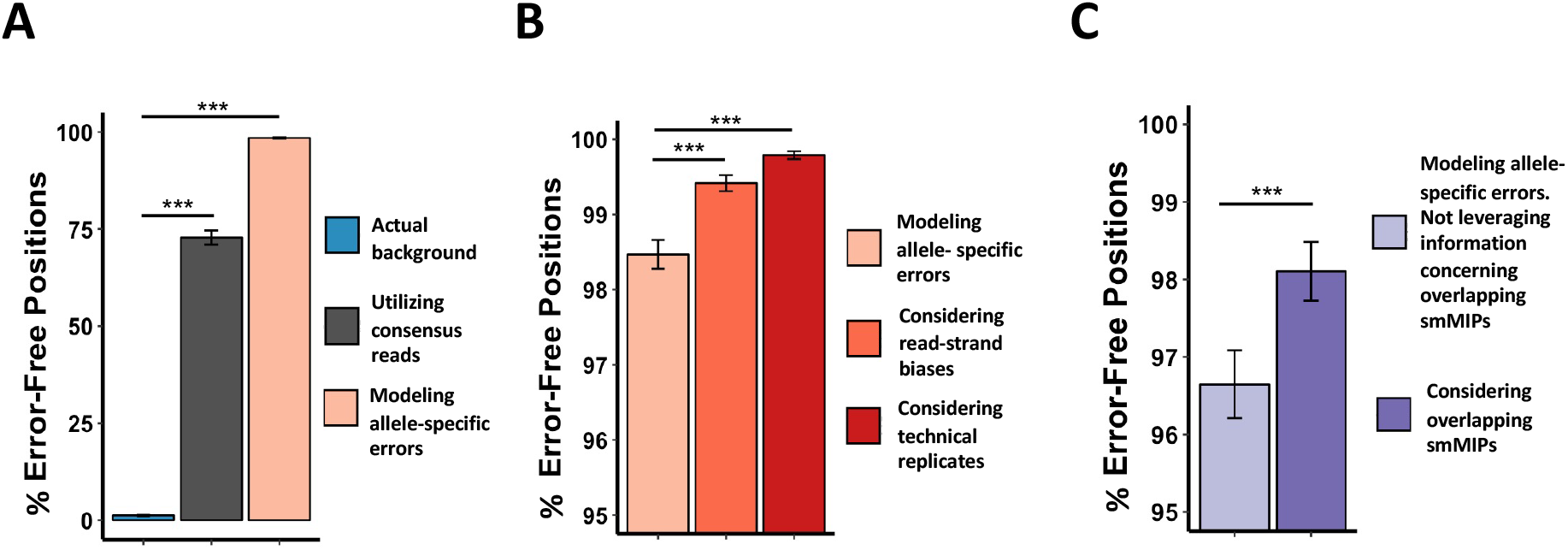
SmMIP-tools uses multiple approaches to supress errors. (A) The percentage of error-free positions across the targeted genomic space before and following error suppression by SSCS assembly or by probabilistic error-rate modeling. The average error background in the cell line DNA mixes is showed for comparison. (B) Percentage of error-free positions following probabilistic error-rate modeling and consideration of information derived either from paired reads or technical replicates. (C) Percentage of error-free positions following probabilistic error-rate modeling alone versus the additional consideration of overlapping smMIPs. Here, only positions covered by overlapping smMIPs were considered to derive the percentage of error-free positions.

Next, we evaluated SmMIP-tools’ performance in differentiating real mutations and errors by using all of the error suppression techniques mentioned above. We required at least 1 SSCS in each strand, and for error rate modelling, we accounted for data derived from separate strands, overlapping smMIPs and technical replicates. SmMIP-tools accurately identified the real mutations among the high background of NGS-associated errors, as evident by a near-perfect trade-off between sensitivity and specificity (Fig. 4A). Sensitivity and precision remained high down to a VAF of 0.005, only decreasing to a lower limit of 9.1% sensitivity and 50% precision for mutations detected in the 0.001<VAF<0.005 range (Fig. 4B).

**Figure. 4.**
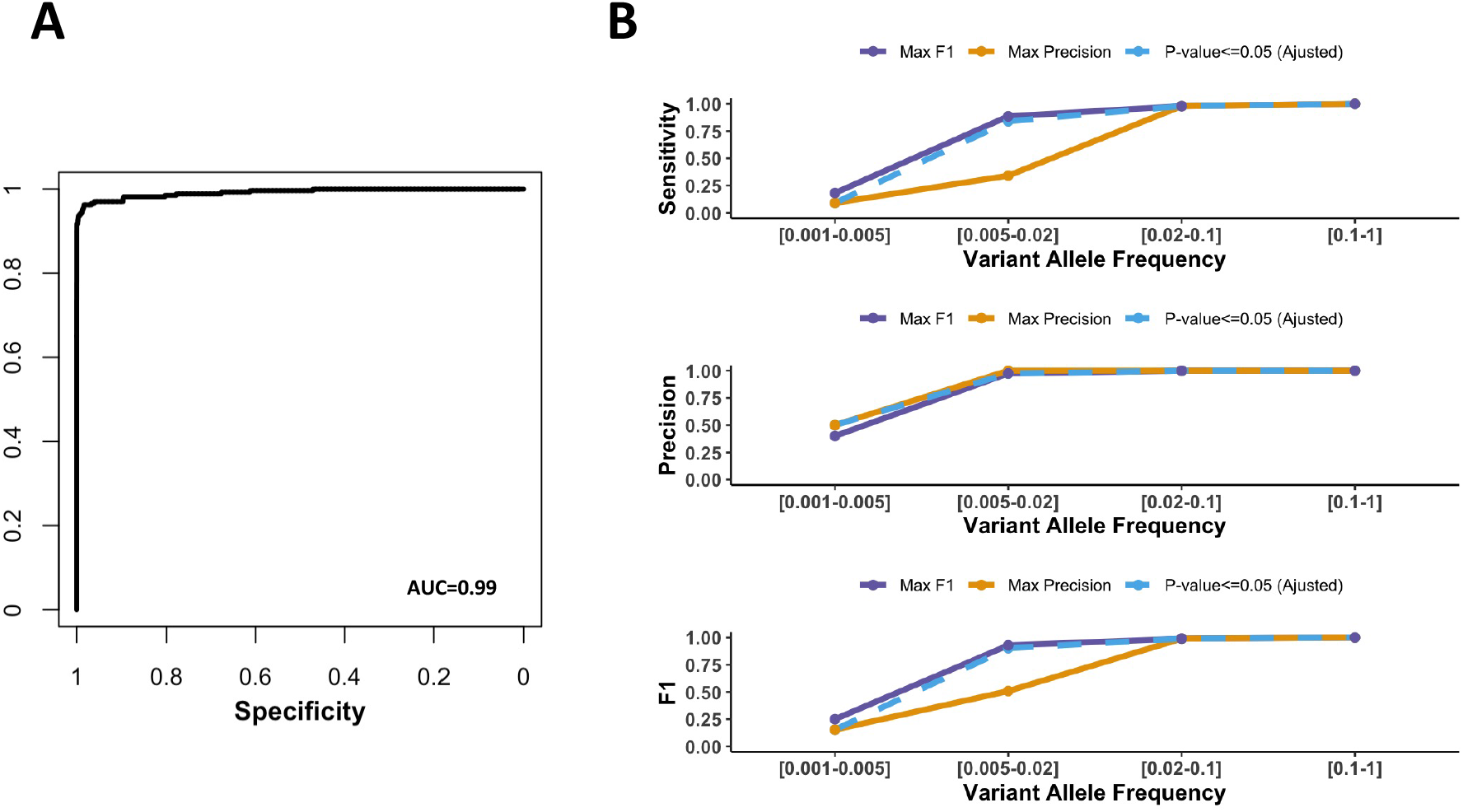
SmMIP-tools accurately differentiates real mutations from NGS-associated errors. (A) Receiver operating characteristic (ROC) curve indicating high performance of SmMIP-tools to detect real mutations in DNA cell line mixes among the high background of NGS-associated errors. Area under the curve (AUC). (B) Accuracy (precision, sensitivity and F1 score) is shown across different VAF ranges. Colored lines represent the results obtained when the core algorithm was set to achieve maximum F1, maximum precision or when the default P-value cut-off (0.05) was used.

Importantly, SmMIP-tools performance using a P-value threshold of 0.05 showed to be remarkably similar to its performance at maximum F1, suggesting that a P-value of 0.05 is a good default cut-off to use when a sequencing cohort does not include artificially diluted samples with known mutations for P-value optimization. Errors are key confounding factors for sensitive detection of low-frequency variants by deep sequencing (Ma et al. 2019). This analysis illustrates SmMIP-tools error-aware variant calling algorithm’s capabilities in accurately detecting mutations with varying VAF ranges among the high number of errors acquired during NGS.

### A comparative analysis between SmMIP-tools output and clinical reports obtained from leukemia patients

As supervised DNA dilutions may be limited in their ability to fully represent clinically relevant mutational processes, we also carried a comparative analysis between SmMIP-tools mutations calling algorithm output and clinical genetic reports generated for patients diagnosed with myeloid malignancies (Methods). Re-sequencing of 168 samples from 162 patients using smMIPs was conducted in technical duplicates. Clinical reports were available for 135 of the patients (Supplemental Table 3). For appropriate comparison with SmMIP-tools, we accounted only for genomic loci covered by both sequencing panels (Supplemental Table 4). Two mutation categories termed “High Confidence” and “Lower confidence” were evaluated based on SmMIP-tools’ generated flags (Methods).

Overall, 95.6% of the somatic SNVs and indels detected using the clinical pipeline were also detected by SmMIP-tools (Supplemental Table 3). Of these, 97.7% are in the “High Confidence” category. Technical issues were identified to be the primary reason for the missing 4.4% of the variants (Supplemental Table 3). On the other hand, SmMIP-tools detected an additional 111 high confidence SNVs (25.2% increase) and 30 indels (32.6% increase) that were not mentioned in the clinical reports of the patients (Fig. 5A, Supplemental Table 5).

**Figure. 5.**
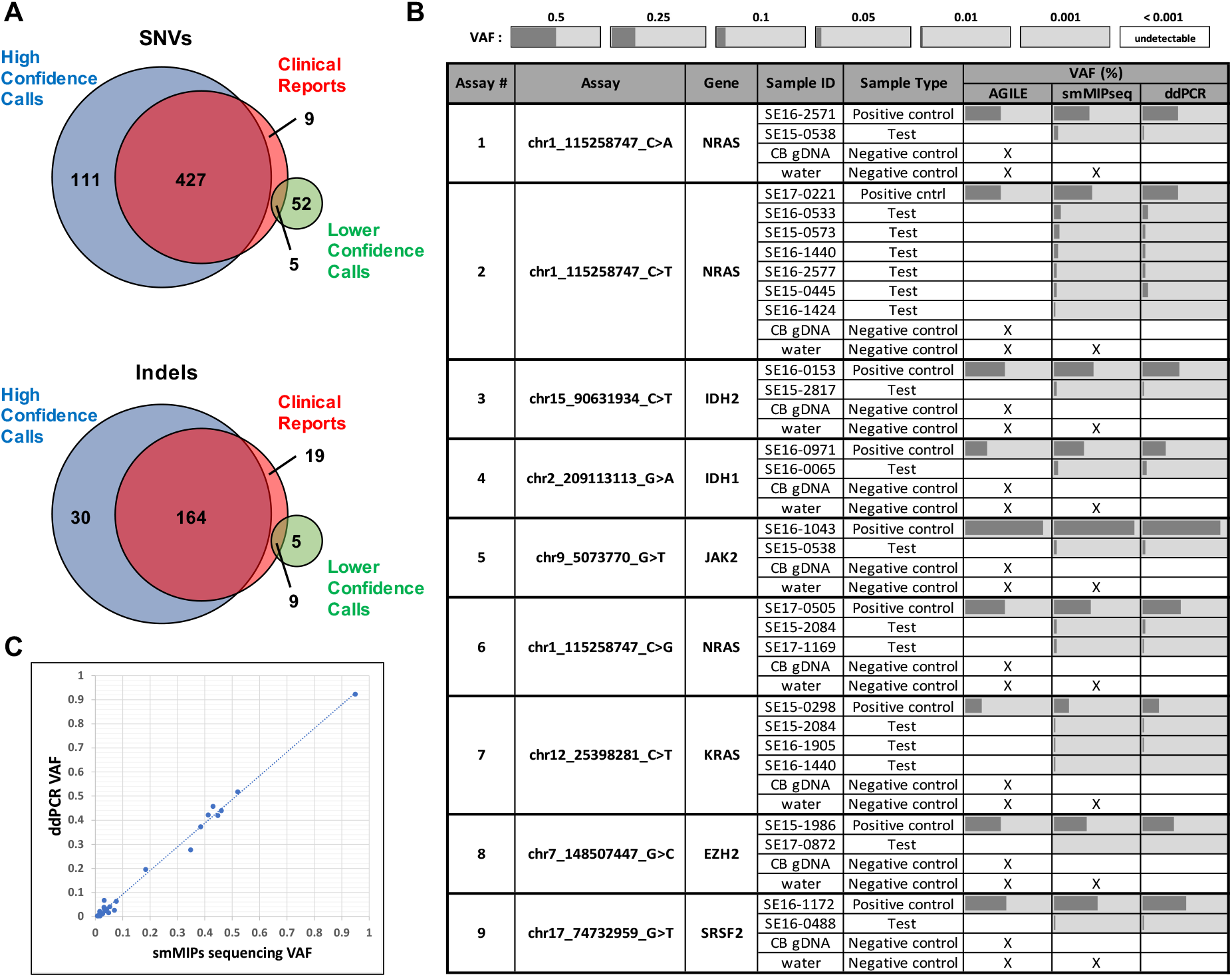
Mutations detected by SmMIP-tools in comparison with patients’ clinical reports. (A) Venn diagrams for SNVs and indels illustrate the number of mutations that are shared between SmMIP-tools output and the clinical genetic reports as well as mutations that differ between the two sources. (B) Validation of a subset of high confidence mutations identified in SmMIP-tools output but not clinical reports by ddPCR. White background indicate that the variant was not reported while light grey background indicates the variant was detected above VAF of 0.1%. Dark grey bars represent the VAFs. X - not tested. (C) Scatterplot showing a strong linear correlation (Pearson r=0.996, P-value=9.81×10^-7^), between VAFs calculated by ddPCR and those derived from the SmMIP-tools output for the 17 high confidence SNVs interrogated in test samples and positive controls.

An average of 29.7 (median=29.6) scaled CADD score (Rentzsch et al. 2019) was calculated for these 111 SNVs, indicating a striking enrichment for deleterious mutations. A subset of high confidence SNVs representing 15.3% of the newly detected SNVs was chosen for validation using digital droplet PCR (ddPCR, Methods). The authenticity of these variants was successfully validated at a 100% rate (Fig. 5B). Furthermore, a high correlation was observed between the VAFs obtained by both smMIP sequencing and ddPCR results (Fig. 5C).

SmMIP-tools was also capable of providing a plausible explanation for the source of some false non-reference alleles, including those detected at relatively high VAF yet not mentioned in the clinical sequencing reports (Supplemental Fig. 3). Information concerning each read’s smMIP-of-origin, reads’ family size, and UMI sequences suggest that these errors could be generated from index misassignment (also termed as “index-hopping”) between multiplexed libraries.

**Supplemental Figure 3.**
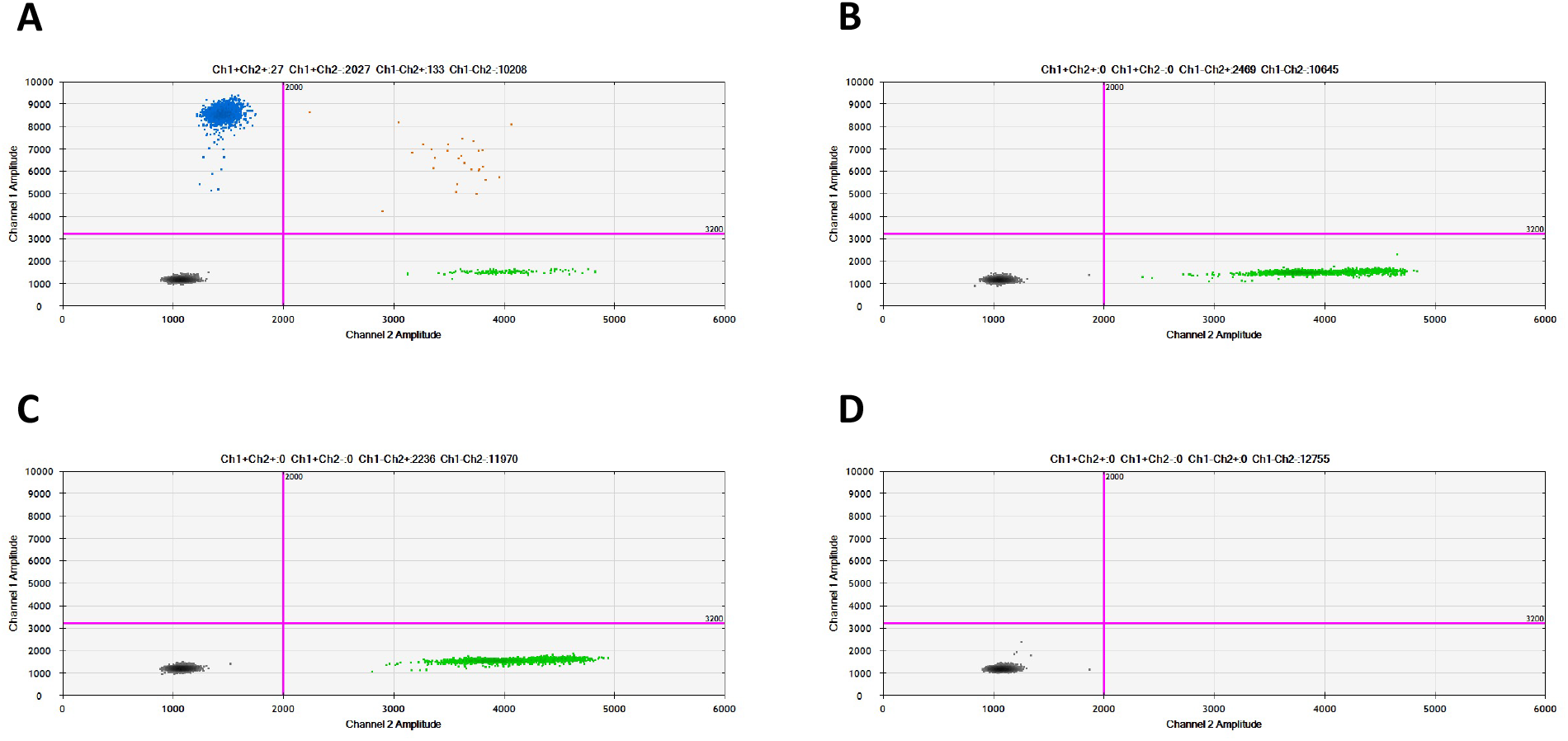
ddPCR results for the oncogenic variant JAK2^V617F^, flagged as “Potential index-hopping”. (A) Sample SE16-015 served a positive control. (B) Inability to detect the mutated allele from the test sample, SE16-0218, where the mutation was detected using smMIP sequencing yet was flagged as an error that may arise due to sample-index misassignment. (C) Cord blood DNA was used as a negative control. (D) A water sample was served as “blank”. Blue dots correspond to the FAM fluorophores that marks the detection of the mutated allele. Green dots correspond to the HEX fluorophores that marks the wild type allele.

Supporting this is the fact that multiple other samples in the cohort presented the same non-reference alleles, some with high VAF (Supplemental Table 5), and shared one sample index barcode with these samples, thus providing an abundant source of reads that could be misassigned. Nine of the 23 variants flagged by SmMIP-tools as “Potential index hopping” were also flagged as “Common SNP” and were detected with atypical VAF far below the expected ~50% for germline mutations. This further supports SmMIP-tools use of UMIs and read-smMIP linkages to flag errors that would otherwise be easily mistaken as true mutations using other pipelines that do not take this information into account

## Discussion

Molecular inversion probes enable cost-effective multiplex targeted sequencing. However, the immense analytical burden associated with handling the resulting data poses a major obstacle to many users of the technology. Several of the fundamental steps in smMIP-derived data interpretation include initial data quality assessment, read processing, alignment of reads, summarizing the raw base calls, and mutation calling. To address the high amount of analytical work associated with NGS data interpretation, many tools have been developed, yet the critical need to address many fundamental analytical steps under a single software suite, especially as they apply to smMIP-derived data, still remains. The need to join together different programs makes the construction of efficient analytic pipelines difficult and time-consuming. Furthermore, and most importantly, some functions that address critical issues specific to the smMIP technology are entirely missing in existing tools. As an example, one key feature that is typically absent from other NGS data analysis tools is the ability to link the reads to their precise probe-of-origin. We have shown that by generating read-smMIP linkages, one can identify and filter chimeric reads and non-informative reads that originated from self-annealing smMIPs. Furthermore, we have shown that read-smMIP linkage can support the suppression of errors by comparing results derived from overlapping smMIPs. We also designed SmMIP-tools to identify the correct position of the smMIP arms’ sequences in the reads’ layout. This unique feature is necessary to prevent mutation calling outside of the target regions of any particular smMIP and support the marking of reads with compromised UMIs to improve the overall quality of the data for downstream analysis.

Defining the absolute ground truth is a major challenge when reporting mutations from NGS data (Koboldt 2020). Errors that arise during library preparation and sequencing are abundant and can easily obscure real mutations. The introduction of molecular barcode sequences into the probes’ sequence (Hiatt et al. 2013) was an essential step in improving the original MIP design towards more accurate mutation calling. In addition to its ability to use UMIs to assemble SSCS, SmMIP-tools provides a versatile error modelling approach to calculate a P-value for every non-reference allele observed in the data to reflect the probability of a false observation. To address different experimental configurations, SmMIP-tools was designed to derive with error rates models using either a single matched control sample, a cohort of controls or with no controls by leveraging data from the experimental cohort itself. Specificity can be improved by modelling all types of base substitutions, insertions and deletions in both the paired reads, overlapping smMIPs, and technical replicates if available.

To our knowledge, SmMIP-tools is the only software that uses sequencing batch information to support variants’ authenticity. In addition to P-values, users are provided with sample-specific and allele-specific information, including the VAF of the called allele in other samples, the number of samples in which the same allele was observed with a higher VAF and the number of other samples in which the allele was detected above the sequencing noise. Other types of outputs that are unique to SmMIP-tools include the number of reads that were grouped to generate individual consensus sequences for every detected non-reference allele and their intra-family VAFs, all of which are important to rank and prioritize mutation calls effectively. We have shown that batch-related UMI information can also be used to flag false positives potentially arising due to sample-index misassignment.

To mitigate index misassignment, specific recommendations, including the use of specific adaptor sequences and sequencing systems have been investigated (Costello et al. 2018). Here we provide a novel *in-silico* approach to identify calls that potentially arose due to index misassignment that could be otherwise mistaken for real mutations using alternative mutation calling algorithms. This functionality of leveraging information other than sample barcode sequences can reduce the cost associated with generating large multiplexed libraries, and thus further increases the smMIP technology’s cost-effectiveness. Nevertheless, it is important to note that unlike our *in-silico* approach, using non-combinatorial dual indexes does allow direct identification and removal of swapped reads which induce sample contamination (Costello et al. 2018).

Sensitive and cost-effective NGS sequencing opens a myriad of clinical applications, including screening of microbial populations, non-invasive prenatal testing and early cancer detection. It can improve routine diagnostic testing of tumours in molecular diagnostic laboratories and inform cancer treatment decisions by enabling longitudinal sequencing efforts to monitor emerging treatment resistant clones. We anticipate that the public availability of SmMIP-tools will lead to improved smMIP-derived data analysis and increased use of the technology.

## Methods

### smMIP panel design to maximize error identification and suppression

Proper smMIP design includes all the discrete genomic elements encased within a single DNA oligonucleotide. This includes a standard backbone sequence, UMIs and custom extension and ligation arms (Fig. 1A). The two halves of the backbone sequence are complementary to sequences located on the forward and reverse PCR primers, thus permitting subsequent PCR amplification of captured sequences (Fig. 1B). We used dual sample indices, each 8 bases in length, on each PCR primer to enable the high multiplex sequencing conducted in this study. We also designed dual UMIs, each 4 bases in length, at each side of the backbone sequence to allow for PCR amplified target sequences to be collapsed to the single smMIP-captured molecule from which they derived. This UMI design allows the detection of up to 65,536 unique molecules per smMIP targeted loci. All the oligos sequences that are needed for successful sequencing are illustrated in Fig. 1.

In this study, we targeted hotspot variants and exons known to be commonly mutated in myeloid malignancies (Supplemental Table 4). smMIPs were designed using MIPgen (Boyle et al. 2014). To maximize the error-correction capabilities of SmMIP-tools we followed two guiding principles. First, the read length of the intended sequencing platform of interest was considered to help determine the optimal target sequence length that will allow every smMIP’s target to be sequenced by both paired-end sequencings reads fully. Second, when a relevant genomic region could not be covered entirely by a single smMIP, we designed additional overlapping smMIPs to tile the region. No preference was set to design overlapping smMIP on opposite DNA strands. Based on analysis of simulated data, smMIPs were designed to align >5 bases apart from the start or end site of their overlapping smMIPs to eliminate erroneous read-smMIP linkages. SmMIPs were ordered as separate standard oligos (Thermo Fisher Scientific), pooled 1:1, deployed on cord blood DNA samples and then empirically rebalanced by increasing the concentration of low-performing smMIPs to achieve a more uniform sequencing coverage across the panel.

### smMIP-Library preparation and sequencing

SmMIP libraries were generated as described elsewhere (Cantsilieris et al. 2017; Hiatt et al. 2013). Phosphorylation of the rebalanced smMIP pool was preformed using a smMIP phosphorylation reaction previously described [18]. Once phosphorylated, the smMIP pool was diluted such that 4ul would achieve a 1:1000 DNA:smMIP molecular ratio in subsequent target capture steps when using 5ul of DNA prepared at 20ng/ul for a total input of 100ng. The probe capture reaction, including the probe hybridization phase, and gap-fill and ligation phase as well as exonuclease treatment, were performed as previously described [8]. PCR amplification of exonuclease-treated smMIP captured products were also performed as previously described [8]. Technical duplicates of PCR amplified libraries were generated for each sample, using unique combinations of barcoded PCR primers. Once generated, sample libraries in each 96-well plate were pooled by volume. Each pool was quantified using Qubit (High-Sensitivity dsDNA Assay, Thermo Fisher Scientific) and underwent a PCR clean-up using MinElute columns (Qiagen). Up to 5ug of input was used per column. Elution was made with 15ul of molecular-grade water. Each pool was subjected to quality control steps, including quantification and fragment analysis on the BioAnalyzer 2100 high-sensitivity assay (Agilent). All the different plate-derived pools were then pooled together by concentration taking into consideration the number of samples in each plate. This final pool then underwent size selection using PippinHT (Sage Science) to mitigate the carryover of smaller and larger products suspected to be self-annealed smMIPs without target sequences and complex oligo hybrids, respectively. Lastly, pooled and size-selected multiplexed smMIP libraries were subjected to sequencing. Bulk cell line libraries were sequenced on the NextSeq using the Mid-Output kit (Illumina). Cord blood, cell line DNA mixes and patients’ libraries were sequenced using a single NovaSeq SP flow cell (Illumina). 10% PhiX was used to ensure sufficient library complexity.

### Derivation of mutation lists for SmMIP-tools benchmarking using cell line DNA mixes

To construct lists of high confidence errors and mutated alleles to benchmark SmMIP-tools mutation calling algorithm, we first sequenced the DNA of 8 cell lines, namely ME-1, Kasumi-1, KG-1A, U937, SU-DHL-10, OCI-AML3, OCI-AML-22, and OCI-AML-8227. Each cancer cell line was sequenced in 4 replicates, and SmMIP-tools was used to report mutations at each of the 32 samples independently using the experimental cohort itself for allele-specific error rate estimations. That is, allele frequencies in each investigated sample were evaluated against those observed in all the samples of the other cell lines. Altogether, we identified 44 high confidence SNVs and 4 insertions (Supplemental Table 2). True positive status was determined if a mutation: 1) Received a P-value<=0.05, 2) Was found in at least 3 out of 4 replicates with VAF>=0.02, and 3) Was supported by at least 1 SSCS from each strand (Read1 and Read2). We removed mutations if they were shared by more than 3 different cell lines to prevent the list from being populated with a large number of common germline mutations. Alleles assigned with a P-value<=0.05 in individual samples but did not pass the additional criteria above were eliminated from benchmarking and were assigned with a status “Uncertain” (Supplemental Table 2). Every other non-reference allele in the remaining interrogated genomic positions was considered as false positive if it was seen in the follow-up sequencing of the 6 DNA mixes (Supplemental Table 2).

For cell line mixes data analysis, a minimum of one SSCS in each strand was required. For the derivation of probabilistic error rate models, a control cohort of 16 umbilical cord blood samples was used. A p-value threshold of 0.05 was set to suppress errors following Bonferroni correction for multiple tests. When calculating the percentage of error-free positions, the focus is on false positives. Therefore, both real mutations and alleles with the status “Uncertain” were removed for this analysis. Alleles with the status “Uncertain” were not included in sensitivity, precision and F1 calculations.

### Generating read-smMIP linkages

To determine the probe-of-origin for every sequenced read pair, SmMIP-tools first searches for smMIPs that their targeted genomic loci, including the extension and ligation arms, substantially overlap with the genomic loci determined by the paired-reads’ alignment to the genome (default 0.95, user defined parameter). Once smMIP candidates are selected, the algorithm proceeds with local alignment of each smMIP’s extension and ligation arms to the reads. The exact probe-of-origin is determined if both of its arms will align the number of UMI bases (here 4nt) from the reads’ extremes. The exact location of UMI bases in each read of the pair (i.e, 5’ or 3’) is determined by the reads’ SAM flags and the number of UMI bases in each read which is automatically determined from the user-provided panel design file obtained from MIPgen (Boyle et al. 2014). When the above alignment expectations are not met, the UMI will be considered unreliable and the paired reads will not be used for any further analysis concerning SSCS.

### Assembly of single strand consensus sequences

SmMIP-tools uses the R-package Rsamtools (Morgan M, Pagès H, Obenchain V, Hayden N. Rsamtools: Binary alignment (BAM), FASTA, variant call (BCF)) to generate base call summaries which are subsequently used to calculate allele frequencies across the target panel. Base call summaries were generated for every UMI-smMIP combination. To minimize information loss, the minimum depth parameter was set to one read, the minimum read mapping quality was set to 50 and the base quality cut-off to 10 (user-defined parameters). At each genomic position, the most frequent allele was determined based on a 70% threshold as previously described (Kennedy et al. 2014) (user-defined parameter). We did not restrict read family size since this information is being captured in the final SmMIP-tools report. The final report allows complete evaluation of the data and is being used to flag and rank reported alleles.

### Probabilistic modeling of error-rates

SmMIP-tools uses the pbinom R function to calculate for each observed allele in a sample of interest a P-value reflecting the probability of obtaining a number equal to or higher than the observed number of non-references supporting reads for the same allele in a single matched control, a larger control cohort or with no controls. If the latter option is chosen, SmMIP-tools uses the entire cohort except for the sample of interest (and its technical replicate, if available) to estimate error rates. Using the cellbaseR R package (Abdallah 2020), sequenced alleles are annotated, and information concerning recurrent cancer mutations and polymorphisms is leveraged to increase sensitivity at those positions. Samples detected with recurrent alleles with VAF>=0.05 (user-defined parameter) were removed from error rate calculations. The allele frequency in all the other samples was set to their median value. To derive binomial probabilities, allele-specific error rates were determined as the sum of all the non-reference supporting reads in all the controls divided by the total number of reads covering the allele. This value was evaluated against the number of non-reference supporting reads and the allele’s coverage in the sample of interest. If any allele was detected by both Read1 and Read2 it received the higher P-value of the two models. This process was repeated for alleles that are covered by overlapping smMIPs and technical replicates. For benchmarking, we modelled allele-specific error rates using a set of 16 control umbilical cord samples where true mutation burden and its effect on real error rates are expected to be low. For comparison with patients’ clinical reports, the experimental cohort itself served as control.

### Mutation categories

By default, SmMIP-tools generates a report which includes baseline mutations calls defined as those mutations detected with a Bonferroni corrected P-value that is lower than or equal to the user-defined P-value cut-off. In this study, we used a cutoff of 0.05. The number of tests used for Bonferroni correction is equal to the total number of alleles that can potentially be detected using the smMIP target panel. These include “A”, “C”, “T”, “G”, “-” (deletion) and “+” (insertion) at every targeted genomic position (Supplemental Table 4). SmMIP-tools uses the R-package cellbaseR (Abdallah 2020) to annotate single base substitutions. To enrich the baseline calls for somatic and pathogenic mutations, we removed common SNPs with a minor allele frequency (MAF)>0.01 as well as mutations assigned with the “potentially germline” flag that was determined based on MAF cut-offs of 0.001<=MAF<0.01 (user-defined parameters). SNVs with the “potentially germline” flag were kept only if they have been reported in COSMIC (Tate et al. 2019). We also removed SNVs that were assigned the “likely benign” flag. Those include mutations with one of the following annotations: “synonymous variant”, “intron variant”, “splice region variant” (rather than “splice acceptor/donor variants”), “non-coding transcript exon variant”, “5 prime UTR variant”, or “3 prime UTR variant”. For Indels, we removed those that fall entirely within intronic regions.

The report generated by the SmMIP-tools mutation calling algorithm also includes additional information concerning consensus reads and details about how baseline called alleles are being represented in other samples in the sequenced cohort. We used this information to derive two categories for mutation ranking, namely “High confidence” and “Lower confidence”.

1) High confidence mutations were determined as those mutations that: a) Received a P-value that is lower than or equal to 0.05. b) Are enriched for somatic and pathogenic mutations as described above. c) Had no flags indicating possible technical issues that may interfere with their detection (i.e., the flags: “Cannot be supported by both Read1 and Read2”, “Cannot be supported by all overlapping smMIPs“, “Cannot be supported by both technical replicates”, “No SSCS support in one of the replicates”, “No SSCS support at all”, “No SSCS support in either Read1 or Read2 in at least one of the replicates”). d) Had no flags indicative of potential false positives (i.e., the flag “Potential index hopping”). e) Did not have the “VAF Warning” flag. The absence of this flag indicates that the mutation was detected with the highest VAF in the reported sample and that its VAF is at least 2-fold (user-defined parameter) higher as compared with all the other samples in the entire cohort. Of note, by default, the algorithm that generates this flag permits that a single sample with a lower VAF will not reach the 2-fold VAF difference. The algorithm was also designed to work properly with authentic recurrent mutations detected as baseline calls in the cohort.

2) Lower confidence mutations were determined as those mutations that received flags indicating potential technical issues that could have interfered with their detection. Those include mutations that were detected only by Read1 or Read2 or only in a single smMIP where overlapping smMIPs were present. Of note, optimal smMIP design where every targeted base is sequenced twice, once with Read1 and once with Read2, can reduce the number of lower confidence calls. Other mutations in this category are mutations that were not detected in both the technical replicates due to low coverage anomalies in one of the replicates. A minimum coverage of 10x from each read (Read1 and Read2) (user-defined parameter) was required to consider the strand-derived information for P-value calculations. For any mutation call, when the coverage does not reach the set threshold, a detailed “Low coverage” flag will be generated. More specifically, we defined and reported lower confidence mutations if: a) They were detected in at least 50% of the available detection options mentioned above (i.e. overlapping sequencing reads, overlapping smMIPs and technical replicates). For example, if a mutation was detected in both Read1 and Read2 but in only one replicate, where overlapping smMIPs do not exist, it will be reported since it was detected in 50% of the possible detection options. If a mutation that is covered by a single smMIP was detected with only Read1 in only one replicate, it would not be reported since it was detected in only 1 of the 4 possible options. b) There were not more than 5 other samples in the entire sequenced cohort where those mutations received a P-value<=0.05. c) There were no other samples that were detected with those mutations with a higher VAF (only relevant to mutations that lack a Cosmic ID).

### Patient cohort

All patients referred to the MPN/leukemia program at the Princess Margaret Cancer Center were approached for targeted sequencing as part of the Advanced Genomics in Leukemia (AGILE) prospective study (Alduaij et al. 2018). All biological samples were collected according to procedures approved by the Research Ethics Board of the University Health Network (01-0573.28) and were viably frozen in the Princess Margaret, Leukemia Tissue Bank. Written informed consent was obtained from all patients in accordance with the declaration of Helsinki.

### Clinical reports

DNA samples extracted from peripheral blood or bone marrow were used for NGS. Sequencing was performed using the TruSight Myeloid Sequencing Panel (Illumina) and run on the MiSeq Illumina platform as previously described using a protocol that was validated by the University Health Network Advanced Molecular Diagnostics Laboratory (Thomas et al. 2017). Briefly, 54 genes implicated in myeloid malignancies were profiled using amplicon-based library preparation (Supplemental Table 4). Data analysis and quality assessment for reporting of SNVs and short indels was performed using NextGene v.2.3.1 (SoftGenetics, State College, PA). Detected variants were then annotated using the WHO classification and diagnostic criteria as previously reported (Barbui et al. 2018) to differentiate between variants of unknown significance and those that are oncogenic. Oncogenic variants detected at coverage >100X, with VAF>5%, were included for subsequent investigation. Known hotspot variants detected below these thresholds were verified using either Sanger sequencing or ddPCR. Detection of recurrent NPM1, insertions or deletions, and FLT3 internal tandem duplications was performed using fluorescent polymerase chain reaction (PCR) followed by fragment analysis. Variants that repeatedly occurred in >10% of cases, excluding mutation in hotspot regions or variants previously reported in the literature, were filtered out.

### Digital droplet PCR

We used ddPCR to validate a subset of SNVs that were detected as “high confidence” mutations by SmMIP-tools but were not reported in the clinical sequencing reports. Nine ddPCR assays were designed to target oncogenic SNVs spanning a VAF range of 0.008-0.077 as determined by SmMIP-tools in 17 patients (n=1 VAF<0.01, n=5 0.01<VAF<0.02, n=8 0.02<VAF<0.05, n=3 0.05<VAF<0.08). Wild type and mutant specific PrimeTime LNA qPCR probes were conjugated to HEX and FAM fluorophores, respectively (Integrated DNA Technologies). 20ng of DNA, originally used for clinical sequencing and re-sequencing using smMIPs, was subjected to ddPCR assays in a 96-well plate according to the manufacturer’s protocol (Bio-Rad). Droplets were generated and read using the QX200 ddPCR system (Bio-Rad) to detect the absolute number of reference (HEX) and non-reference (FAM) alleles. VAF was calculated as the number of FAM captured targets over the total number of targets (FAM+HEX). As previously reported (Shlush et al. 2017), variants were considered positive if >3 droplets were captured in the mutant (FAM) channel with a final VAF>0.001. Samples with the same mutations detected by both the clinical pipeline and SmMIP-tools were used as positive controls. Cord blood DNA was used as a negative control and water was used as a blank. ddPCR assay conditions and associated primer and probe sequences are supplemented (Supplemental Table 6).

## Supplementary information

**Supplemental Figure 1.** Addressing issues related to the smMIP technology by linking reads to their probe-of-origin with read-smMIP linkages

**Supplemental Figure 2.** Simulating smMIPs to assess the performance of read-smMIP linkages generation

**Supplemental Figure 3.** ddPCR results for the oncogenic variant JAK2^V617F^, flagged as “Potential index-hopping”

**Supplemental Table 1.** Comparison between SmMIP-tools and popular analytical methods for NGS

**Supplemental Table 2.** True cell lines mutations and alleles that were removed from the benchmarking process

**Supplemental Table 3.** Sequenced samples and clinical genetic results

**Supplemental Table 4.** Targeted genomic loci

**Supplemental Table 5.** SmMIP-tools mutation calls in the clinical cohort

**Supplemental Table 6.** Details concerning ddPCR assays preformed

* Supplemental tables are provided as separated files.

## Acknowledgements

This project was supported by work done at the Advanced Molecular Diagnostic Lab (AMDL) at the Princess Margaret Cancer Centre. The authors also thank the team at the Princess Margaret Genomics Centre for their sequencing services and genome informatics at the Ontario Institute for Cancer Research for providing high-performance computing. We thank Dr. Jean Wang, Dr. Qiang Liu, Dr. Eric Lechman and Dr. Héléna Boutze for providing us with the cell lines used for SmMIP-tools benchmarking.

## Authors’ contributions

JJFM optimized the design of smMIPs, generated NGS smMIP libraries, preformed ddPCR assays, contributed to data analysis and wrote the manuscript. JJCC managed clinical sequencing and provided the clinical genomic reports. LIS and JED contributed to panel optimization and sequencing. MDM and AA managed patients’ selection, and provided the clinical genomic reports and DNA samples from the Princess Margaret, Oncology and Hematology Tissue Bank for smMIP resequencing. SA wrote the code and the manuscript, contributed to data analysis and directed and supervised all aspects of the study. All authors read and approved the final manuscript.

## Funding

The work conducted in the SA lab is supported by the Investigator Award received from the Ontario Institute for Cancer Research with funds from the province of Ontario and by the University of Toronto’s Medicine by Design initiative, with funds from the Canada First Research Excellence Fund (CFREF). JJFM is personally funded by the Canadian Institutes of Health Research Doctoral Award: Frederick Banting and Charles Best Canada Graduate Scholarships (FBD-170928).

## Competing interests

LIS serves as a consultant for Sequentify LTD and Metasight LTD. All other authors declare that they have no competing interests.

## Availability of data and materials

The source code for SmMIP-tools, its manual and additional scripts used for data generation and analysis are all available on github (https://github.com/abelson-lab/smMIP-tools). Raw sequencing data generated in this study have been submitted to the European Genome-Phenome Archive (EGA; https://ega-archive.org) under accession number EGAS00001005359. All other information concerning the use of SmMIP-tools, the benchmarking process and the comparison of SmMIP-tools results with the patient’s clinical reports are included in the method section and supplemental files.

## Notes

### Competing Interest Statement

Liran I. Shlush serves as a consultant for Sequentify LTD and Metasight LTD.

### Summary of Updates

Supplemental tables added. All other aspects remain unchanged.

